# Diffusion Acceleration with Gaussian process Estimated Reconstruction (DAGER)

**DOI:** 10.1101/436550

**Authors:** Wenchuan Wu, Peter J Koopmans, Jesper Andersson, Karla L Miller

## Abstract

**Purpose:** Image acceleration provides multiple benefits to diffusion MRI (dMRI), with in-plane acceleration reducing distortion and slice-wise acceleration increasing the number of directions that can be acquired in a given scan time. However, as acceleration factors increase, the reconstruction problem becomes ill-conditioned, particularly when using both in-plane acceleration and simultaneous multi-slice (SMS) imaging. In this work, we develop a novel reconstruction method for in-vivo MRI acquisition that provides acceleration beyond what conventional techniques can achieve.

**Theory and Methods:** We propose to constrain the reconstruction in the spatial (k) domain by incorporating information from the angular (q) domain. This approach exploits smoothness of the signal in q-space using Gaussian processes, as has previously been exploited in post-reconstruction analysis. We demonstrate in-plane acceleration exceeding the theoretical parallel imaging limits, and SMS combined with in-plane acceleration at a total factor of 12. This reconstruction is cast within a Bayesian framework that incorporates estimation of smoothness hyper-parameters, with no need for manual tuning.

**Results:** Simulations and in vivo results demonstrate superior performance of the proposed method compared with conventional parallel imaging methods. These improvements are achieved without loss of spatial or angular resolution and require only a minor modification to standard pulse sequences.

**Conclusion:** The proposed method provides improvements over existing methods for diffusion acceleration, particularly for high SMS acceleration with in-plane undersampling.

## 1. Introduction

As neuroscience studies aim for higher spatial and angular resolution, image acceleration is an increasingly important part of the diffusion MRI (dMRI) toolkit. The acquisition time of a dMRI sequence is determined by the number of diffusion volumes (q-space samples) and the scan time for each volume (in k space). The number of diffusion directions is an important determinant of data quality for dMRI methods that benefit from high angular resolution (e.g. for tractography). Diffusion MRI data are typically acquired using 2D echo planar imaging (EPI). While EPI is itself quite time efficient, dMRI sequences are overall very inefficient because ≥50% of the sequence time is typically dedicated to the diffusion preparation. This inefficiency is compounded by the fact that each slice is acquired sequentially, such that the scan time per volume is proportional to the number of slices. For fixed coverage, higher spatial resolution requires more slices and therefore increased scan time. When scan time is limited, there is a fundamental trade-off between spatial coverage, spatial resolution and angular resolution (density of q-space sampling).

In recent years, acquisition and reconstruction methods have been developed to mitigate this trade-off. One major improvement is simultaneous multi-slice (SMS) imaging (1–3) which can reduce the volume acquisition time, enabling more diffusion directions within the same scan time. SMS techniques have particularly benefitted from improved k-space encoding schemes like blipped-CAIPI (3). However, due to the intrinsically low SNR of dMRI data and the ill-conditioning at higher multiband (MB) factor, slice acceleration at 4× or above is still difficult to achieve. Moreover, SMS accelerations rely on the same coil hardware as is used to achieve in-plane acceleration (4,5), which is crucial at high spatial resolution to reduce EPI distortion and blurring. The combination of in-plane and SMS accelerations is thus highly desirable, but currently ill conditioned. Regularized parallel imaging methods could potentially alleviate this problem by enforcing some prior information in the spatial domain (6–10). Although undersampling factors above 10 have been reported in structural or fMRI, high acceleration is considerably more challenging in dMRI. To take one prominent example, the Human Connectome Project exhaustively evaluated SMS accelerations for both fMRI and dMRI. The optimized HCP protocol utilized MB=3 (without in-plane undersampling) for dMRI, compared to MB=8 for fMRI (11).

An alternate approach is to accelerate in q-space. Compressed sensing (CS) has been used to improve the estimation of diffusion profiles using a small number of q-space samples and estimating the diffusion signal at unseen locations in q-space (12–17). These approaches exploit the information redundancy in q-space and impose a sparsity constraint under a data transform (e.g. wavelet, total variation) or in a simple model space (e.g. summation of tensors).

Finally, a series of methods have proposed joint k-q acceleration, with data undersampling in both k space and q space (18–21). These methods all cast joint k-q acceleration as a compressed sensing problem and differ primarily on the specific sparsifying transform or model that is used. Most of these k-q methods focus on in-plane undersampling to enable acquisitions in comparable scan times to single-shot acquisitions but at much lower distortion for a given resolution.

In this work, we propose a new method to accelerate dMRI acquisition by incorporating sharable information between q-space samples. One property that has been exploited for acceleration is the smoothness of signal in q-space (20). Gaussian Processes (GPs) (22) offer a general framework for smoothing and interpolation of signals in a model-independent fashion. GPs use a covariance function to capture the observed statistical dependences of the observed data. In diffusion imaging this has been used to utilize the smoothness in q-space to correct for eddy-current induced distortions and subject movement (23,24) and replace outliers (25).

Building on these approaches, we have integrated GPs into joint k-q reconstruction to achieve high acceleration in the context of both in-plane and slice-wise acceleration. We call this approach Diffusion Acceleration with Gaussian process Estimated Reconstruction (DAGER). We use a Bayesian framework that learns the smoothness hyper-parameters from the data itself, with no need for manually tuning reconstruction parameters (22). We demonstrate this approach using both numerical simulations based on retrospectively under-sampled data and prospective in vivo acquisitions. Two acceleration schemes are evaluated: a very high in-plane undersampling factor (exceeding the theoretical limit of conventional parallel imaging) and using a high SMS acceleration factor with in-plane undersampling (the primary intended application). Both simulations and in vivo results demonstrate superior performance of DAGER compared with conventional parallel imaging methods. These improvements are achieved without loss of spatial or angular resolution and require only a minor modification to standard pulse sequences.

## 2. Theory

### 2.1 GP modelling of dMRI signal

A GP is a statistical model defined as a probability distribution over functions where function values evaluated at any arbitrary set of input points have a joint Gaussian distribution (22). Diffusion MRI signal can be seen as the output of a real-valued function *S*(**q**_i_). The input **q**_i_ is specified by a diffusion encoding direction and a b value. If we model the function *S* using GPs, then any finite collection of dMRI signals **u** = *S*(**q**) at an input set 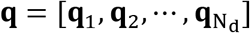 follows a multivariate Gaussian distribution:

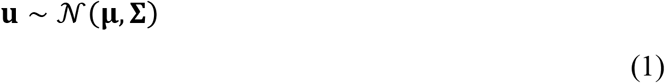

This distribution is fully specified by a mean vector **μ** (N_d_×1) and a covariance matrix **Σ**(N_d_×N_d_). **u**(N_d_×1) contains dMRI signal from a single voxel for all diffusion volumes. N_d_ is the number of diffusion volumes. The spatial distribution of signals (i.e. covariance between voxels) is temporarily ignored for simplicity of notation, which will be covered in Section 2.3.

An important feature of Gaussian processes is that the covariance is modelled by a covariance function *C*, which is typically characterised by a small number of hyper-parameters **Θ**. A typical property motivating the choice of covariance function is that the covariance between two variables is higher when the input points are closer to each other. This means that one must be able to define a distance measure in the space of the independent variables. Once the covariance function is determined, the elements of the covariance matrix can be calculated accordingly:

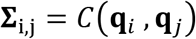

Diffusion MRI signal is typically acquired on one or several q-space shells, each corresponding to one b-value. A reasonable choice of covariance function can be defined based on the angular difference between diffusion directions and the distances between shells, although equivalent functions could also be defined for other q-space sampling schemes, such as Cartesian sampling in diffusion spectrum imaging. The spherical covariance function (26) has been shown to have good agreement with the covariance observed from real dMRI data (24). In the current work, we focus on single-shell acquisition (a single b-value), with the corresponding spherical covariance function having a single parameter *a*, and being given by:

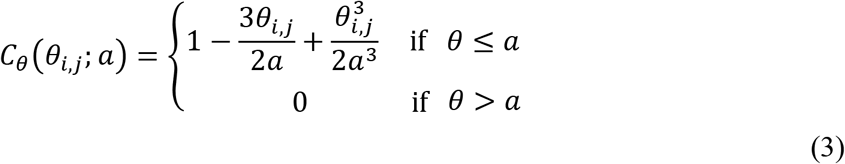

Here, *θ*_*i,j*_ denotes the angle between the two vectors pointing from the q-space origin to the qspace points **q**_*i*_ and **q**_*j*_, defined as *θ*_*i,j*_ = cos^−1^ 〈**q**_*i*_, **q**_*j*_〉 A smaller angle *θ* is associated with a larger covariance between the two q-space points. The decay rate of the covariance with respect to *θ* is governed by the smoothness hyper-parameter *a*, which also serves as an angular threshold for covariance.

To account for the variance of signal and noise in real dMRI data, two other hyper-parameters, λ and σ, are also included in the covariance function: *C*(**q**_*i*_, **q**_*j*_) = λ*C*_*θ*_(*θ*_*i,j*_;*a*) + σ^2^ δ_*i,j*_, where λ captures the variability of the signal and σ^2^ represents the variance of the noise that is uncorrelated in q-space. *δ*_*i,j*_ is the Kronecker delta function. The covariance matrix can be derived accordingly

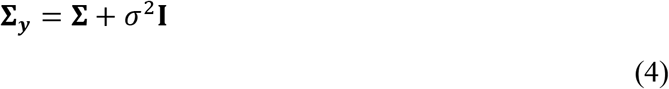

with element of **Σ** defined as *Σ*_*i,j*_ = λ*C*_*θ*_(*θ*_*i,j*_;*a*), capturing the q-space smoothness, and **I** is an identity matrix.

If prior information about the mean signal for each diffusion direction is available, the mean function can also be modelled explicitly (22). Otherwise, the mean of a GP model is usually set to zero. Note a zero-mean prior is not a strict limitation, as the posterior mean is not constrained to zero.

### 2.2 GP prediction of dMRI signal

GP methods allow us to place a prior distribution directly on the functions, which is transformed to a posterior distribution after having been fit to data. This posterior can be used to make prediction of unseen dMRI signal. This has been previously used as a data analysis approach for removing distortion and discarding outliers (23,25). In the context of image reconstruction, it can be used to fill in missing data in order to accelerate acquisition. We begin by recapping this method, which will then adapt for use in k-q image reconstruction.

Assume we have a set of dMRI images 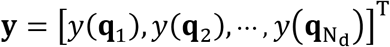, which contain additive noise:

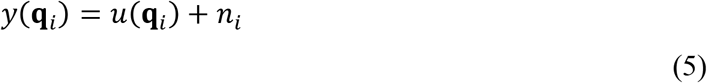

The noise *n*_*i*_ is assumed to follow a2n independent and identically distributed (i.i.d.) Gaussian with zero mean and a variance of 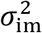 The likelihood for this data is then:

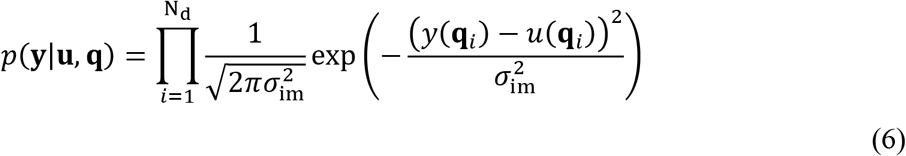

If we cast our estimation of **u** in terms of Bayes’ theorem with a GP prior (Eq. 1), the maximum a posteriori (MAP) estimation provides a point estimate (27):

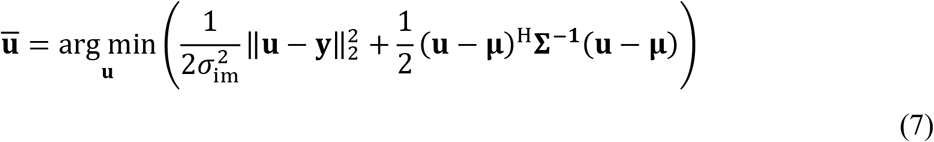

whose analytical solution is given by 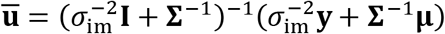, which is a linear combination of prior and data with weights determined by the prior covariance and noise variance.

### 2.3 dMRI reconstruction with a GP prior constraint

The previous sections have described GPs in terms of a single voxel, i.e., signal that is localized in image space. In image reconstruction, data are acquired in k space and our goal is to estimate the unknown signal in image space. The dMRI measurement can be described as a linear model:

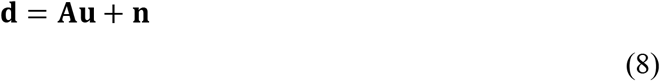

where **u**(N_v_N_d_×1) is extended to contain the signals for all diffusion volumes from all voxels. N_v_ is the number of voxels in each volume. **d**(N_v_N_d_N_c_×1) denotes the acquired k-space data for all N_d_ diffusion volumes. Note the vector **y**in the previous section is the observation in image space. N_c_ is the number of coils. **A**(N_v_N_d_N_c_× N_v_N_d_) is the system matrix combining sensitivity encoding, Fourier transform and the k-space sampling operation. **n**(N_v_N_d_N_c_×1) is the measurement noise, which is assumed to follow an i.i.d. Gaussian distribution with a zero mean and variance 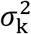.

We now extend the GP model (Eq. 1) to include the spatial distribution of signals

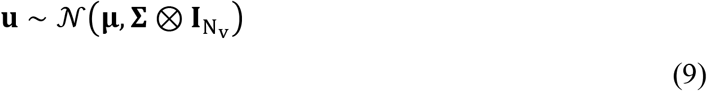

where **μ** and **Σ** have the size of N_v_N_d_×1 and N_d_×N_d_, respectively. 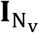 is a N_v_×N_v_ identity matrix ⊗. is the Kronecker product, which is used to generate the full covariance matrix (N_v_N_d_×N_v_N_d_):

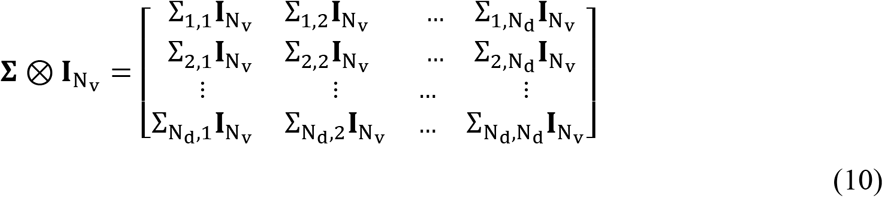

The covariance values are purely a function of angular distance between directions (i.e. the same function as described above). Although each pair of directions may have a unique covariance value Σ_i,j_, there is only one set of hyper-parameters to estimate.

Conventional parallel imaging aims to find the maximum likelihood estimate (i.e. the point estimate that does not include any prior constraints) by solving a minimization problem (28):

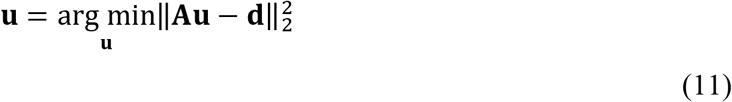

By incorporating a GP prior on the signal, the MAP estimate can be obtained as (27):

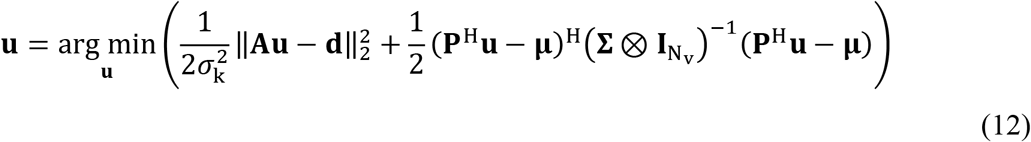

Disregarding the **P** matrix (discussed below), this is a straightforward extension of the MAP estimation given in Eq. 7, which extends the maximum likelihood to include a second term enforcing the prior. This MAP formulation is written to resemble standard linear image reconstructions with a regularisation between two cost functions. The unusual feature here, however, is that the regularization parameter (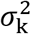 and **Σ**) is clearly defined based on the Bayesian likelihood, as discussed in the next section.

**P** denotes the motion-induced phase errors and is the conjugate transpose operator. The phase errors are primarily driven by bulk and cardiac motion during the diffusion preparation and have no relation to the q-space locations. The acquired complex signal thus violates the GP assumption that information is correlated in local q-space. The term **P**^H^ is included in the reconstruction to realign the image phase across all diffusion volumes. The phase errors can be measured from a low-resolution navigator (29,30) or estimated from the imaging data itself (31) provided conventional parallel imaging reconstruction is able to eliminate most aliasing artifacts.

### 2.4 GP hyper-parameter estimation

Solving Eq. 12 requires knowledge of the hyper-parameters **Θ** = [λ, a, σ] encapsulated in thecovariance matrix **Σ** Using Bayes theorem, the optimal hyper-parameters can be estimated from image data by maximising the marginal likelihood *p*(**y|q, Θ**). In practice, one typically minimises the negation of the log of the marginal likelihood

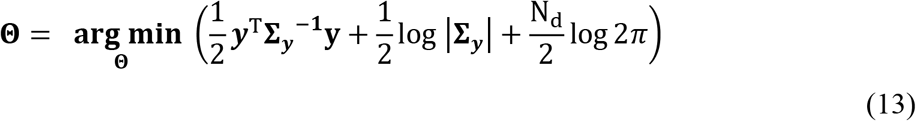

where |**Σ**_**y**_| is the determinant of **Σ**_**y**_. In this work, an iterative scheme is used to estimate the optimal hyper-parameters.

## 3. Methods

### 3.1 Image reconstruction and hyper-parameter estimation

An iterative scheme is used to jointly estimate the optimal hyper-parameters and the underlying images: in each iteration, a set of hyper-parameters is calculated (Eq. 13) using the images reconstructed from the previous iteration; then a new image reconstruction is conducted with the updated hyper-parameters (Eq. 12). As the iteration goes on, it is expected that both the hyper-parameter estimation and the image reconstruction can be improved.

The noise variance estimated from the reconstructed images 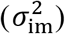 is different from the noise variance of the k-space data 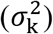, thus one cannot directly use 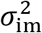 in the MAP reconstruction (Eq. 12). Nevertheless, the propagation of noise in parallel imaging can be quantitatively measured based on sensitivity maps (5), which provides a way to derive σ_k_ σ_im_.

The pipeline of the proposed method is summarised below:

**Initial**: Conventional SENSE is used to reconstruct all diffusion volumes, which produces an image initialization **u**^0^.

The noise variance of the k-space data is estimated from 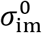 in 1000 randomly chosen brain voxels as:

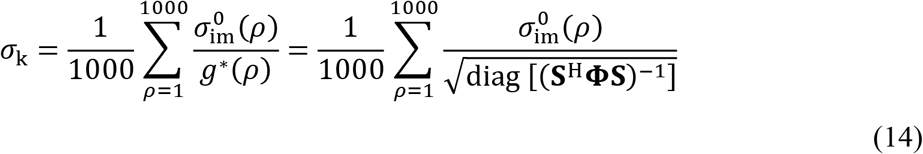

where *g*^*^ the noise propagation function, derived from the coil sensitivities **S** (N_c_ × 1000) and **Φ**(N_c_ × N_c_) is the coil noise covariance matrix (5). While the hyper-parameter σ_im_ is updated each iteration, σ_k_ is calculated in the first iteration and fixed thereafter.

**Iteration n**: Each iteration first updates the hyper-parameters and then the image reconstruction based on the outputs of the previous iteration.

The optimal hyper-parameters 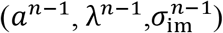 are estimated from the signal across all diffusion directions extracted for 1000 randomly chosen brain voxels in **u**^n−1^. The final estimates are an average of the estimates from all voxels. A covariance matrix **Σ**^*n*−1^ is then constructed and the posterior mean **u̅**^n−1^ is calculated according to Eq. 7.

Finally, we reconstruct all diffusion volumes using **u̅**^n−1^ and **Σ**^n−1^ as prior constraints as the solution to the optimization:

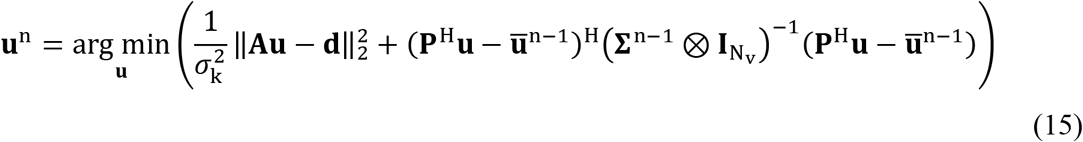

**Output**: We stop iterating when a fixed number of iterations is reached or the normalised ║**u**^n^ − **u**^n−1^║/║**u**^n−1^║ is below a pre-set threshold. The last reconstruction (Eq.15) will be the final result.

### 3.2 k-q sampling

For joint k-q reconstruction, it has been reported that improved reconstructions can be achieved by using different k-space sampling patterns for different diffusion volumes (19–21). Smoothness in q-space means that nearby q-space points share more common features (i.e. the images are more similar than distant points, Fig. 1a). We explore the extent to which we can improve the DAGER joint reconstruction by using different k-space undersampling patterns within a local neighborhood in q-space. Specifically, we use a graph model approach to design the proposed k-q undersampling.

**Figure 1.**
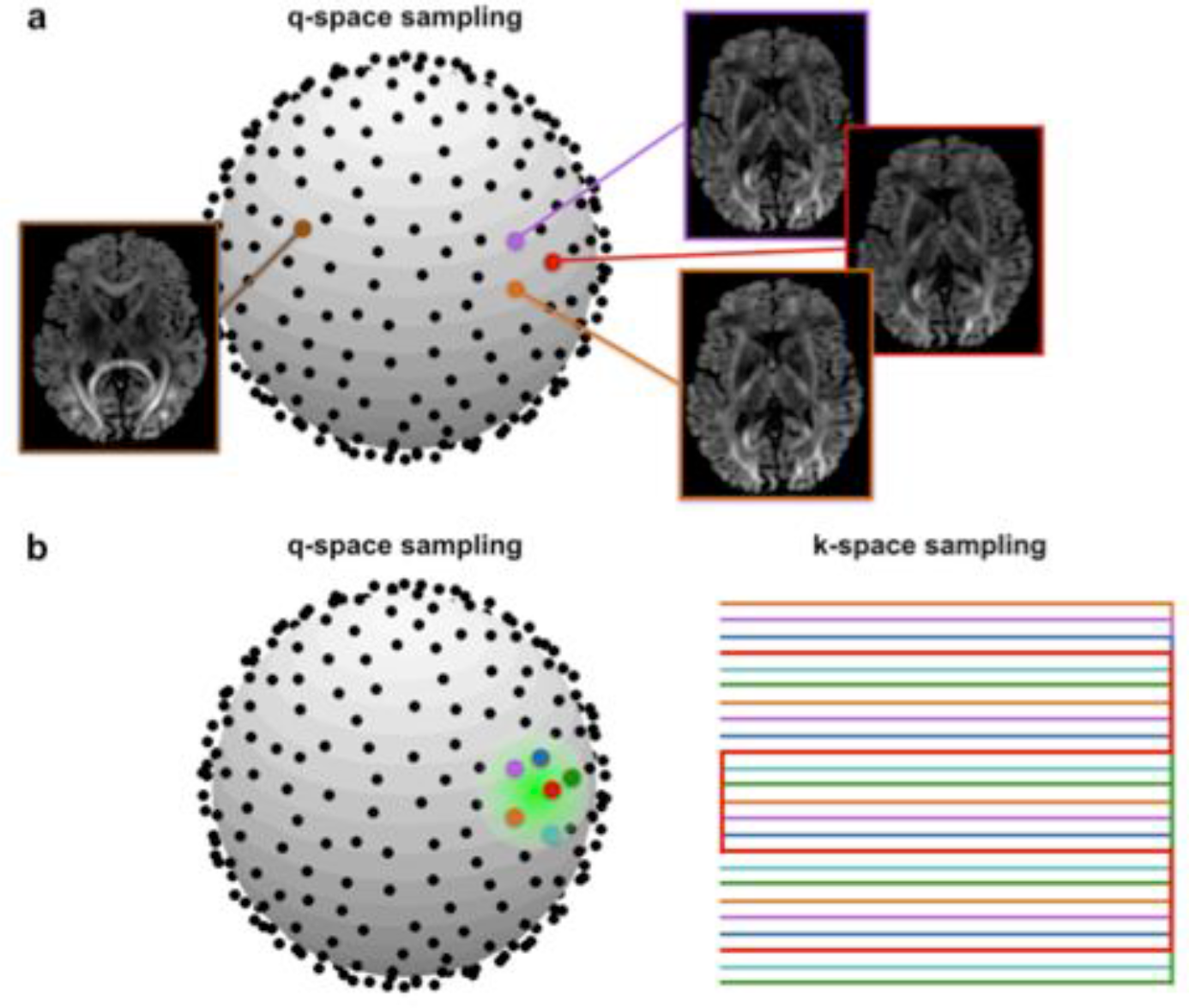
a). DAGER reconstruction takes advantage of local smoothness in q-space: dMRI images that are near to each other in q-space look very similar. b). The proposed k-q sampling scheme: using different k-space sampling patterns in the local q-space neighborhood. Each color corresponds to a different sampling pattern in k-space as shown on the right.

For EPI with an undersampling factor of R, there exist R unique k-space sampling patterns that can be applied at different q-space points. These patterns are produced by shifting the under-sampled trajectory along the phase-encoding direction and slice direction (for SMS acquisition). For each q-space location, a neighbourhood is defined including itself and its R-1 nearest neighbours in q-space (measured by angular distance). The task of assigning different k-space sampling patterns for the q-space points within a local neighbourhood (Fig. 1b) can be equivalently described as a graph-colouring problem. In this formulation, each q-space point corresponds to a vertex and an edge between two vertexes exists if the corresponding two q-space points are within the same neighbourhood. If the constructed graph can be coloured using R different colours while obeying the constraint that no connected vertexes share the same colour, the resultant colouring scheme provides the desired k-q undersampling. Graph colouring is an “NP complete” problem, which means no efficient algorithms are available to provide the exact optimal solution. However, it also establishes an equivalence to all NP-complete problems, for which a number of efficient algorithms exist. In this work, we applied a greedy algorithm to approximate the solution (32).

### 3.3 Simulation

One common way to validate an acceleration method is to acquire fully sampled k-space, which is then retrospectively under-sampled to generate simulation data sets. However, this approach is not very compatible with dMRI acquisition, as single-shot EPI is heavily smoothed due to the rapid 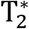 decay, while multi-shot EPI suffers from shot-to-shot phase variation caused by subject motion. Instead, our validation uses a simulation derived from a simplified signal model that was fit voxel-wise to high-quality, in-vivo data. This is particularly helpful for evaluating the performance of DAGER with and without phase corruption. However, this simplified representation of dMRI signal is by construction smooth in q-space and is not perfectly representative of in-vivo data. These simulations thus represent a well-conditioned data environment, and ultimately it is crucial to evaluate performance on in-vivo data.

Simulations were constructed based on dMRI data from the Human Connectome Project (HCP), consisting of three shells (1000, 2000 and 3000 s/mm^2^) and 90 diffusion directions per shell, which had been pre-processed to account for eddy currents and distortions (33). The data were fit with a ball-and-stick model (34,35), which was used to generate simulated data based on the same model. We simulated in-plane undersampling with a small number of channels, aimed to test the limits of the proposed method by using extreme acceleration along one dimension. This included the extreme case where the number of channels was exceeded by the undersampling factor, for which the information supporting the reconstruction would have to come from q-space, demonstrating the central DAGER concept. We also conducted SMS simulations targeting the actual application we would like to achieve in practice, which was to improve the ability to accelerate in-plane and through-plane simultaneously. Therefore, the same number of channels were used for SMS simulation as for the in-vivo data acquisition (following section).

#### 3.3.1 In-plane undersampling

Diffusion MRI data sets with 128 directions and b=1000 s/mm^2^were simulated. Multi-channel data were generated with coil sensitivity maps measured from an 8-channel head-coil (https://www.ismrm.org/mri_unbound/simulated.htm). Complex random noise was added to the data. Three in-plane undersampling factors R= 4,6,10 were evaluated, with R=10 exceeding the theoretical parallel imaging limits for the simulated 8-channel coil. Simulations were generated with and without acquisition-to-acquisition phase errors. The phase errors consisted of a constant offset randomly selected between [−π, π] and a linear phase term in image space, corresponding to a shift along k_y_ by a random distance between [0, 2Δk_y_] where Δk_y_ is the phase encoding step size. The effects of non-linear phase errors are demonstrated in the experiments performed in vivo.

#### 3.3.2 SMS with in-plane undersampling

An SMS data set with four-fold acceleration (MB=4) was simulated from four slices of single-band image data with 192 directions, b=1000 s/mm^2^ and 31.5mm slice gap. Coil sensitivity maps were measured using a 32-channel coil. The blipped-CAIPI (3,36) scheme was used to introduce a FOV/4 shift between adjacent SMS slices. In-plane undersampling factor R=3 was applied, leading to a total acceleration factor of 12. Noise and phase errors were added to the data as described in the previous section.

### 3.4 In vivo experiments

Four subjects were scanned on a Siemens 7T scanner using a custom pulse sequence. Informed consent in accordance with local ethics was obtained before each scan. SMS data was acquired in all four subjects, while additional in-plane undersampling scans were performed on one subject.

#### 3.4.1 Sequence implementation

A 2D spin-echo dMRI sequence was modified to include SMS acquisition and the blipped-CAIPI encoding scheme (3). After each imaging echo, a 2D navigator can optionally be acquired following a second refocusing pulse. Two k-q sampling strategies were implemented: ‘fixed’ sampling with the same k-space lines acquired at each q-space location, and ‘variable’ sampling using the graph-coloring algorithm described above to sample different k-space locations in a local q-space neighborhood. Coil sensitivities were measured using the FLEET-ACS method (37,38) to provide more robust ACS data in the presence of motion and respiration. For in-plane undersampling acquisitions, a bipolar diffusion preparation (twice-refocused spin echo (39)) scheme was used to reduce eddy current effects and a 64×64 navigator was acquired following each imaging echo.

A major challenge for SMS diffusion imaging at 7T is specific absorption rate (SAR). Three aspects of the sequence implementation were optimized to reduce SAR for SMS acquisitions. First, RF pulses were designed to be low SAR. Excitation used a conventional MB pulse and refocusing used a MultiPINS pulse (40), both with TBWP = 2.52 and duration = 10.9ms. Second, the diffusion module used a modified monopolar preparation (single 180° pulse), in which part of the second diffusion gradient is negated and placed before the 180° pulse to reduce eddy currents. Finally, it was found that the initial SENSE reconstruction was able to provide a good estimation of the phase errors, enabling us to eliminate the navigator for SMS acquisitions, thereby removing one RF refocusing pulse. The final sequence for the MB=4 acquisition had low SAR level (<50% of scanner limits).

#### 3.4.2 In-plane undersampling

Diffusion MRI data were acquired from subject 1 using a 32-channel coil. Two acceleration factors R=3 and R=6 were acquired with 1.5×1.5×1.5 mm^3^ resolution (matrix size 168×168, 40 slices, b=1000s/mm^2^ and 128 directions, TR/TE=6500ms/82ms, total scan time 13.8min). The R=3 data were acquired with partial Fourier 6/8, which were used as reference for comparison to R=6 data.

Navigator and b=0 data were reconstructed using conventional SENSE, which might incur significant artifact at high acceleration factors. Hence, for the R=6 imaging data, the navigator was under-sampled by R=4 to reduce artifacts while producing similar distortions as the imaging data; the b=0 data were acquired using multi-shot EPI (6 segments). We acquire five b=0 volumes, one of which had reversed phase encode direction to enable correction of susceptibility induced distortion.

#### 3.4.3 SMS with in-plane undersampling

SMS data were acquired in four subjects. Sets of four slices (MB=4) were acquired simultaneously using the blipped-CAIPI scheme, where a FOV/4 shift was introduced between adjacent SMS slices. 21 sets of MB slices with 31.5mm slice gap were acquired, resulting in a total of 84 slices (126mm) for full brain coverage. In-plane undersampling R=3 was also applied, for a total undersampling factor of 12. Other scan parameters were 1.5 mm^3^ isotropic resolution, matrix size 156×156, TR/TE=3500ms/71ms, b=1000s/mm^2^, 192 diffusion directions, total scan time 11mins. Four sets of b=0 data were acquired using multi-shot EPI (12 segments), including one volume with reversed phase-encoding direction for distortion correction.

Single-band (SB) data, with each slice acquired separately, were acquired from all subjects for comparison to SMS data. For the subjects 1-3, a set of single-band data was acquired with the same FOV (i.e. 84 slices) as the SMS protocol, which was used as a full FOV reference. For these scans, a longer TR (12s) and a smaller number of diffusion directions (i.e. 50) were used to achieve similar acquisition time. For subject 4, three repetitions of single-band data were acquired using the same sequence parameters as in the SMS protocol but containing only 21 slices to match the TR, leading to a total acquisition time of 33min. This data was used to provide a high-SNR reference to identify the degree of angular blurring incurred by the DAGER reconstruction.

### 3.5 Reconstruction implementation

Images were reconstructed using both DAGER and conventional parallel imaging methods, specifically SENSE for in-plane parallel imaging and SMS-SENSE for slice acceleration, which were implemented with custom code.

The SMS-SENSE reconstruction implemented in the work follows a 3D k-space formulation (41,42) where in-plane under-sampling and SMS acquisition are treated as under-sampling along k_y_ and k_z_ dimension. A 2D-SENSE (43,44) method is used to reconstruct the data. Due to the ill-conditioning of the reconstruction problem, Tikhonov regularization is used in SENSE and SMS-SENSE reconstruction (41). It is not trivial to find the optimal regularization parameter. Inappropriate choice of regularization parameters may lead to noisy reconstruction (too small regularization) or aliasing (too large regularization). In the current work, we use the L-curve approach (45) to determine the regularization parameter (Supplementary Fig. S1) and the optimal regularization parameter (corner of L-curve) is used for SENSE and SMS-SENSE results.

The implementation details of DAGER reconstruction are as follows:

#### 3.5.1 Phase error estimation

In simulation, the phase was estimated using the central k-space (32×32) of the imaging data. Based on the simulated motion (zeroth and first-order phase), this approach provides perfect knowledge of phase errors.

For the non-SMS in vivo data, for which a navigator was acquired, a conventional SENSE calculation was applied to reconstruct the navigator image. This was then smoothed by a 2D hamming filter to suppress ringing artifacts and noise. The phase of the image was used as an estimate of the phase errors. We refer to this approach as “navigator based” due to the use of an explicitly acquired navigator.

For SMS in vivo data, which we did not acquire a navigator, the phase errors were estimated from the imaging data itself (31). Images were first reconstructed using SMS-SENSE, followed by a total variation de-noising (46). In this work, we found the reconstruction with the optimal regularization parameter still contained some residual aliasing artifacts. As minimal aliasing is crucial for DAGER to achieve accurate phase estimate, a lower regularization parameter is used in this reconstruction (Supplementary Fig. S1). The reconstructed image was used as an initialization for DAGER reconstruction. The phase of the de-noised image was used as the phase-error estimate. We refer to this approach as “navigator free” since the phase correction does not require an explicitly acquired navigator.

#### 3.5.2 DAGER reconstruction

The symmetric property of the diffusion signal was considered in simulation, where the covariance function was calculated with the angle *θ* defined as *θ*_*i,j*_ = cos^−1^|〈**q**_*i*_, **q**_*j*_〉| (24). For in vivo data, the signal symmetry was not incorporated in order to reduce the effects of distortion induced by eddy currents and/or Maxwell terms on DAGER reconstruction. This led to a definition of *θ*_*i,j*_ = 〈**q**_*i*_, **q**_*j*_〉.

Hyper-parameters were calculated by solving Eq. 13 using the conjugate gradient method. To avoid possible negative values for signal variance and noise variance, the model was modified such that *e*^λ^ and *e*^σ^ were estimated rather than λ and σ: *C*(**q**_*i*_, **q**_*j*_) = *e*^λ^*C*_*θ*_(*θ*_*i,j*_;*a*) + *e*^2σ^δ_*i,j*_.

Iteration stopped when either the maximum iteration number (set to 20) was reached or the normalized solution update (║**u**^n^ − **u**^n−1^║/║**u**^n−1^║) was below 0.002.

### 3.6 Analysis

#### 3.6.1 Simulation data

For the simulation data, normalized root-mean-squared-error (NRMSE) values were calculated between the reconstructed images and the noise-free ground-truth with a brain mask to emphasize the reconstruction errors within the brain region. This metric was used to compare different acquisition and reconstruction options under the condition of ground truth.

#### 3.6.2 Tensor and tractography modeling

The in vivo data were first processed using Topup and Eddy (23) to correct image distortions. These data were then fitted with a tensor model using DTIFIT (47) and, separately, a Bayesian ball-and-stick model with two fiber populations within a single voxel (BEDPOSTX) (34,35). Probabilistic tractography of major white matter tracts was conducted using the AutoPtx tool in FSL (https://fsl.fmrib.ox.ac.uk/fsl/fslwiki/AutoPtx) (48). AutoPtx defines 14 major white matter pathways, represented by 27 sets of masks in the right and left hemispheres in a pre-defined atlas that avoids any operator bias. Tractography was not applied to the single-band data from subject 4 due to the limited field of view.

#### 3.6.3 Investigation of induced angular blurring

DAGER uses a GP-based smoothness prior on q-space, and as such it has the potential to induce angular blurring if it overestimates the smoothness. To evaluate the accuracy of fiber orientation, we compared the fiber orientations between the SMS images reconstructed using DAGER and SENSE and a high-SNR, non-SMS reference in subject 4. The reference data were constructed by averaging two repetitions single-band data (SB-2ave). For all datasets, the mean of the posterior distribution of the ball-and-stick fiber orientations estimated was used for comparison. A mask was used to exclude fibers having a wide orientation distribution around the principle diffusion direction, which was generated from the ‘dispersion’ map (BEDPOSTX output) using a threshold of 0.03. The angular differences for the two fiber populations were calculated between the SMS images and SB-2ave, as well as between an independent set of single-band data (SB-1ave) and SB-2ave.

#### 3.6.3 Investigation of number of directions

In order to use smoothness in q-space to improve k-space reconstruction, there must be nearby q-space samples that lie within the extent of this smoothness. We investigated the effect of the number of diffusion directions on the different reconstructions. The 192-direction data set was sub-sampled to achieve approximately uniform q-space coverage with 160, 128 and 64 directions. Finally, we investigated the effect of error in the estimation of the smoothness hyper-parameter using the set of 64-directions. Two additional SMS-DAGER reconstructions were applied with the hyper-parameter a fixed to 1 (to include ~12 points in each q-space neighborhood) and π (to use all q samples) throughout the reconstruction, where the estimated value for a is 0.7 (having ~7 points in each q-space neighborhood).

Simulation and acquisition parameters, reconstruction algorithms and data analysis described above are summarised in Table 1.

**Table 1.** Summary of simulation and acquisition parameters, and reconstruction algorithms and analysis methods used in data processing.

|  | Acquisition |  |  |  |  |  |  |  | Reconstruction |  |  |  | Analysis |  |  |  |
| --- | --- | --- | --- | --- | --- | --- | --- | --- | --- | --- | --- | --- | --- | --- | --- | --- |
|  | Subj. ID | Diff. dir. | b value (s/mm<sup>2</sup>) | MB/ R | TR/TE (s/ms) | TA (min) | Nav. | No. of slices | Ave. | No. of coils | SENSE | SMS-SENSE | DAGER | DTI FIT | BEDPOSTX | AutoPtx |
| In vivo protocols |  |  |  |  |  |  |  |  |  |  |  |  |  |  |  |  |
| SMS with in-plane undersampling | Subj. 1 2 3 4 | 192 | 1000 | 4/3 | 3.5/71 | 11 | - | 21 slice group | 1 | 32 | - | Y | Y | Y | Y | Y |
| Single-band reference | Subj. 1 2 3 | 50 | 1000 | -/3 | 12/71 | 11 | - | 84 | 1 | 32 | Y | - | - | Y | Y | Y |
|  | Subj. 4 | 192 | 1000 | -/3 | 3.5/71 | 33 | - | 21 | 3 | 32 | Y | - | - | Y | Y | - |
| In-plane undersampling | Subj. 1 | 128 | 1000 | -/3,6 | 6.5/82 | 13.8 | Y | 40 | 1 | 32 | Y | - | Y | Y | Y | - |
| Simulation protocols |  |  |  |  |  |  |  |  |  |  |  |  |  |  |  |  |
| SMS with in-plane undersampling | Subj. 1 | 192 | 1000 | 4/3 | 5.5/89 | - | Y | 1 slice group | 1 | 32 | - | Y | Y | - | - | - |
| In-plane undersampling | Subj. 1 | 128 | 1000 | -/ 4, 6, 10 | 5.5/89 | - | Y | 1 | 1 | 8 | Y | - | Y | - | - | - |

## 4. Results

### 4.1 Simulations of in-plane acceleration

Fig. 2a shows the reconstruction results of simulated data with in-plane undersampling factors of 4, 6 and 10 using the variable k-q sampling. The ground truth is shown on the left. Due to the simulation of only eight coil elements, conventional SENSE cannot reconstruct the images correctly for any undersampling factor. The reconstruction is substantially improved using the DAGER method, which reduces the NRMSE by a factor of ~9 for the R=10 acceleration compared with the SENSE reconstruction, with residuals that appear noise-like rather than structured. Both the variable k-q sampling scheme and phase-error correction play important roles in the reconstruction, as demonstrated in Supplementary Fig. S2.

**Figure 2.**
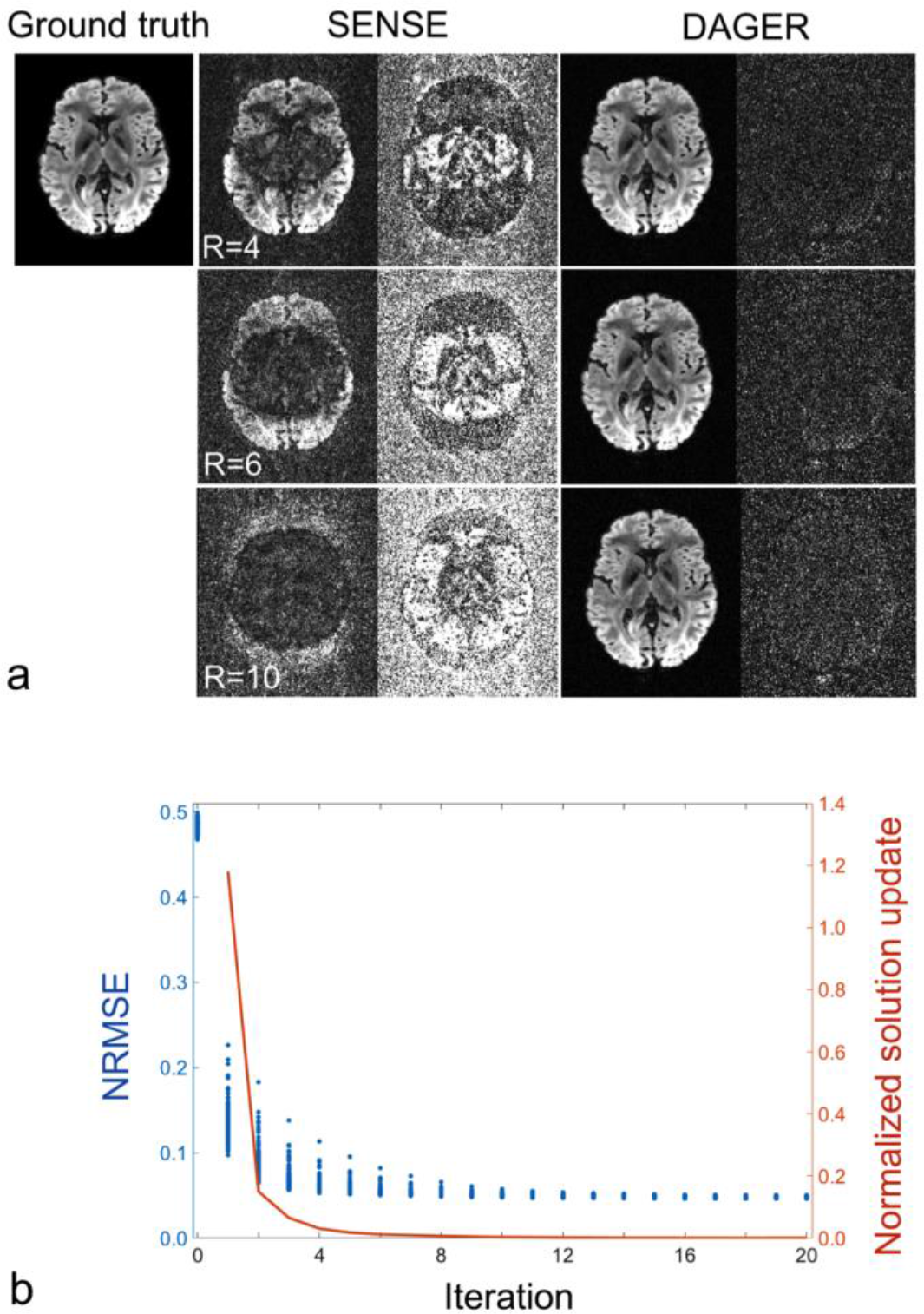
Simulation of in-plane undersampling. a) Three undersampling factors R=4, 6 and 10 are evaluated. Reconstructed images are shown on the left for each method. Difference from ground truth is shown on the right for each method, scaled by a factor of 3 for SENSE and 10 for DAGER. NRMSE values for SENSE are 0.36, 0.48 and 0.63 and for DAGER are 0.04, 0.05 and 0.06 (for R=4, 6, 10 respectively). Note the simulations with R=10 exceed the theoretical limit of conventional parallel imaging, but that nevertheless DAGER reconstruction manages to recover the underlying image with good fidelity. b) Convergence of the DAGER reconstruction for R=6. Blue: NRMSE for all diffusion directions over 20 iterations (each point represents one diffusion direction). Red: Normalized solution update ║**u**^n^ − **u**^n−1^║/║**u**^n−1^║ in 20 iterations. Note the iteration-0 corresponds to the initial SENSE reconstruction.

Fig. 2b shows the convergence of the DAGER reconstruction with R=6 data. The NRMSEs for all diffusion volumes decrease monotonically with iteration and reach a minimum value after about 13 iterations. Although NRMSE is a useful metric to evaluate potential stopping criteria, ground truth is not available in practice and we must rely on metrics like the normalized solution update. The normalized solution update has similar decay behaviour as the NRMSE, which can be used to monitor the reconstruction progress. After 15 iterations, the image update is below 0.002 and the iteration is stopped.

DAGER reconstruction at different noise levels demonstrate that image fidelity (as reflected in NRMSE) is not very sensitive to the estimated value of σ_k_ for SNR≥ 20 (Supplementary Fig. S3). Moreover, σ_k_ is accurately estimated and reconstructed NRMSE are close to optimal at SNR≥ 10, which is crucial given that this parameter is not iteratively updated.

### 4.2 Simulations of SMS with in-plane undersampling

DAGER reconstructions of simulated SMS data are shown in Fig. 3. SMS-DAGER achieves a considerably improved reconstruction compared with SMS-SENSE (tailored SENSE reconstruction for SMS data (41)), reducing NRMSE by factors of 5 - 6. Navigator-based and navigator-free phase corrections achieve comparable fidelity of reconstruction, with slightly higher error in the navigator-free reconstruction (see inset values in Fig. 3). The strong artifacts in the navigator-free SMS-DAGER reconstruction are mostly located in the background where phase estimates are purely noise. The quality of navigator-free reconstruction within the brain suggests the initial SENSE reconstruction used to estimate the phase errors provides a good estimation of the phase errors (for brain region).

**Figure 3.**
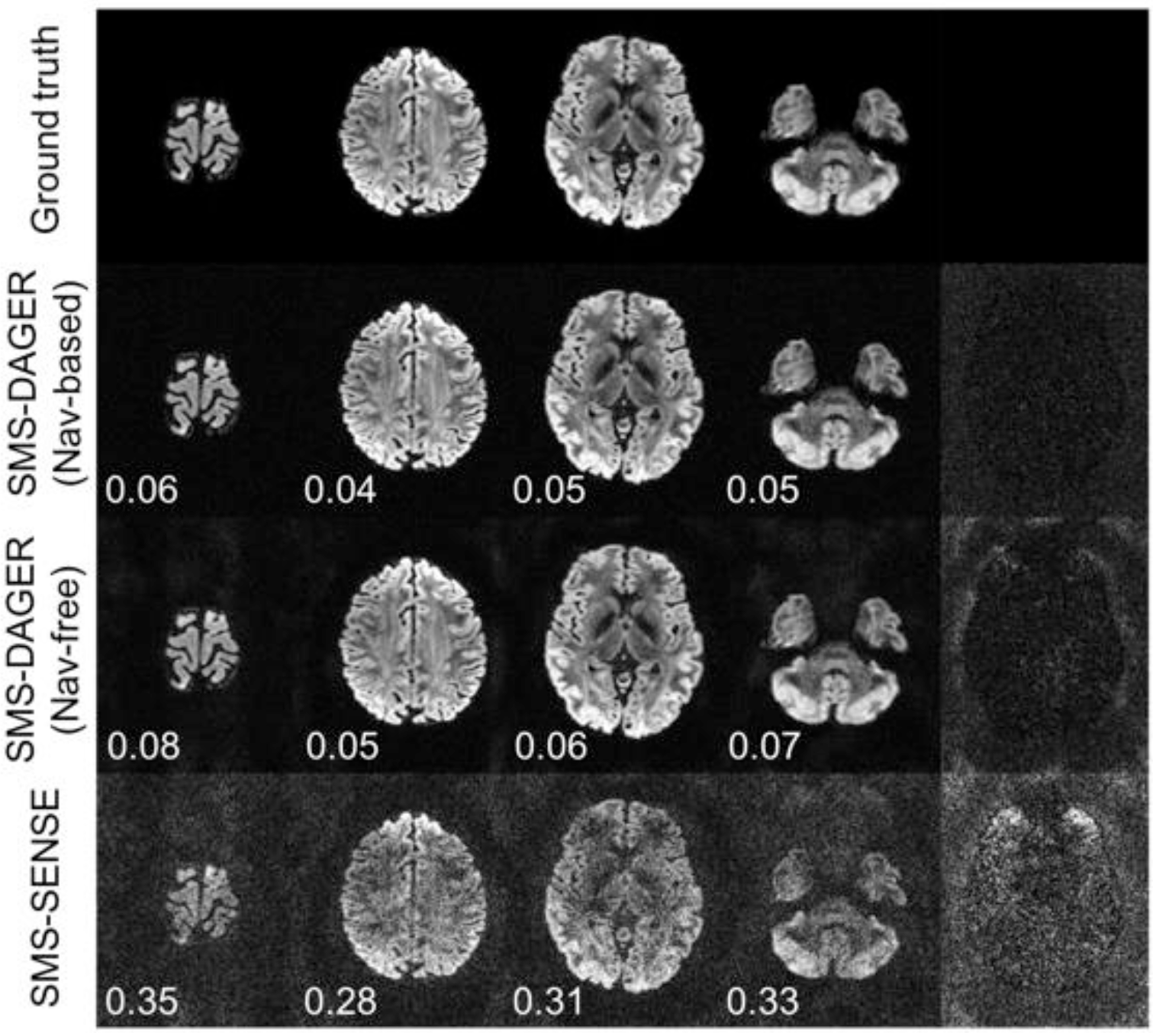
Simulation of SMS data with total acceleration of 12 (MB=4 and in-plane R=3). Three reconstruction approaches are evaluated: SMS-SENSE, SMS-DAGER with navigator acquisition (Nav-based) and SMS-DAGER without navigator acquisition (Nav-free). The four columns on the left are reconstructed images for the four simultaneously-excited slices, with NRMSE values given on the bottom left (calculated for the brain region only). The rightmost column contains the difference images for the third slice, which are scaled by a factor of 5 for SMS-SENSE and 10 for SMS-DAGER.

### 4.3 In-vivo in-plane acceleration

Supplementary Fig. S4 shows in-plane acceleration results from subject 1. Conventional parallel imaging reconstruction at R=6 suffers from noise amplification and residual aliasing even with a large number of coil elements (32ch). DAGER reconstruction is considerably improved. As in simulation, phase-error correction with and without an explicitly acquired phase navigator demonstrate very similar performance. Therefore, the navigator-free method was adopted for the rest of this work to avoid the SAR penalty associated with an additional refocusing pulse for the navigator. Compared with the R=3 image, the R=6 reconstruction with DAGER demonstrates reduced image distortion (visible in the frontal lobes) and improved delineation of structural boundaries.

One indirect way to characterize dMRI data quality is through the estimation of secondary fiber populations within a voxel, where ARD (automatic relevance determination) is used for deciding if a secondary fiber is supported by the data (35). Detection of multiple fibers requires both high contrast-to-noise ratio (CNR) and high angular resolution. As shown in Fig. 4, conventional parallel imaging at R=6 fails to capture many secondary fiber populations, most likely due to the low CNR. In this same data, secondary fibers are recovered throughout the brain in the DAGER reconstruction. This indicates that not only is the CNR high in the DAGER reconstruction, but there is not excessive angular blurring. The reference SENSE reconstruction of R=3 data provides a good CNR with negligible residual artifacts.

**Figure 4.**
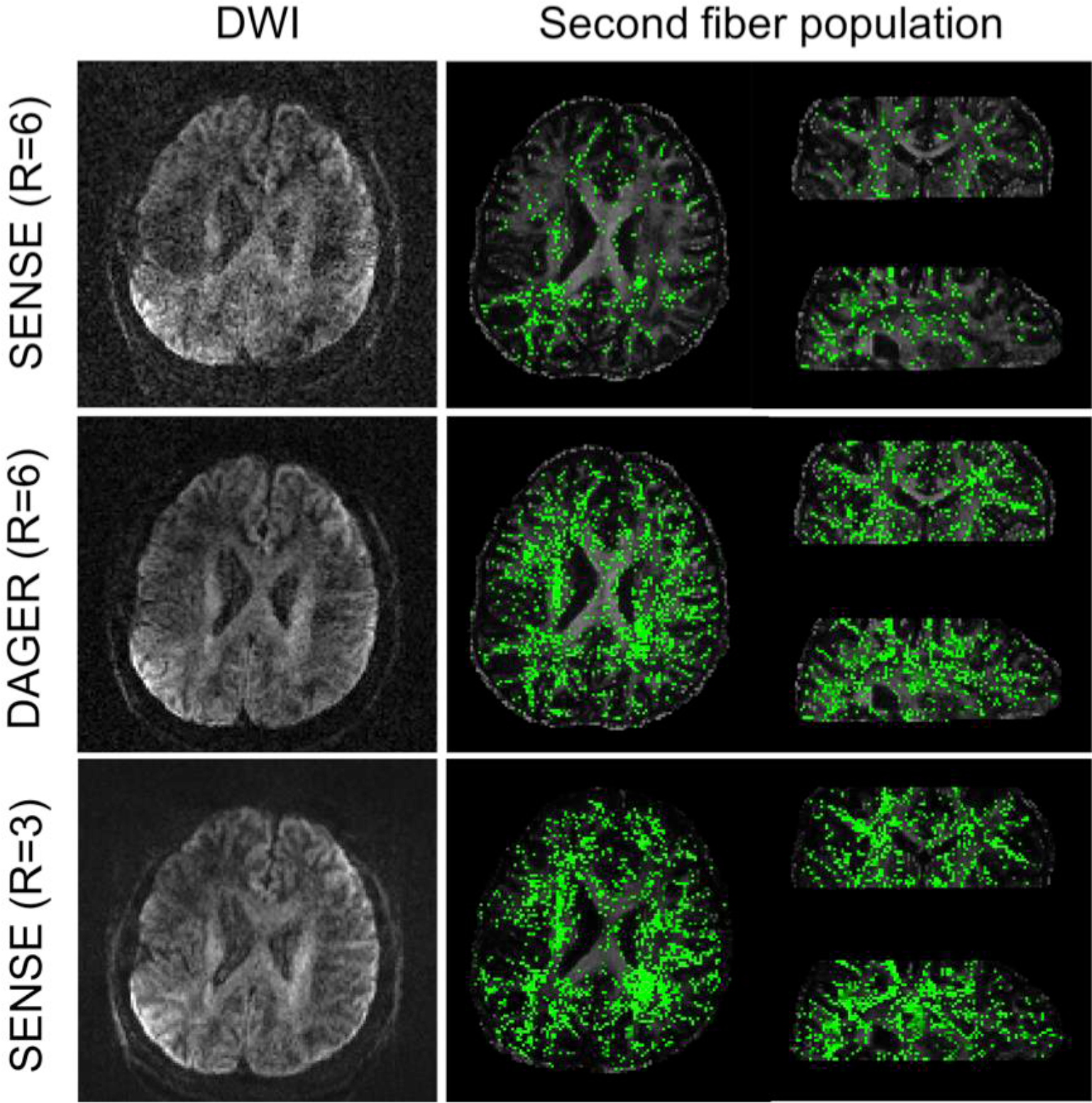
Multiple fiber populations within a single voxel are estimated from R=6 data reconstructed with SENSE (top) and DAGER (middle), and R=3 data reconstructed with SENSE (bottom). The R=3 data are used as a reference here. The second fiber populations for R=3 data, DAGER (R=6) data and SENSE (R=6) data are 89355, 78867 and 23645, respectively. The same threshold (0.05) is used for all data sets.

### 4.4 In-vivo SMS with in-plane acceleration

Fig. 5 shows the reconstruction of SMS data acquired in vivo using both SENSE and DAGER reconstructions. Conventional SENSE reconstruction suffers from amplified noise due to the high acceleration applied in both through-plane and in-plane. From the same data, the DAGER method manages to reconstruct high-quality images with suppressed noise and aliasing artifacts. As shown in Supplementary Fig. S5, the SMS images reconstructed with the DAGER method demonstrate consistent contrast with the single-band reference, which is an average of three repetitions. In addition, the noise level of the SMS image is similar to that of the single-band reference despite a three-fold reduction in scan time, suggesting GP prior can effectively aid in eliminating noise by exploiting the common features between diffusion volumes (or equivalently, by adaptive smoothing).

**Figure 5.**
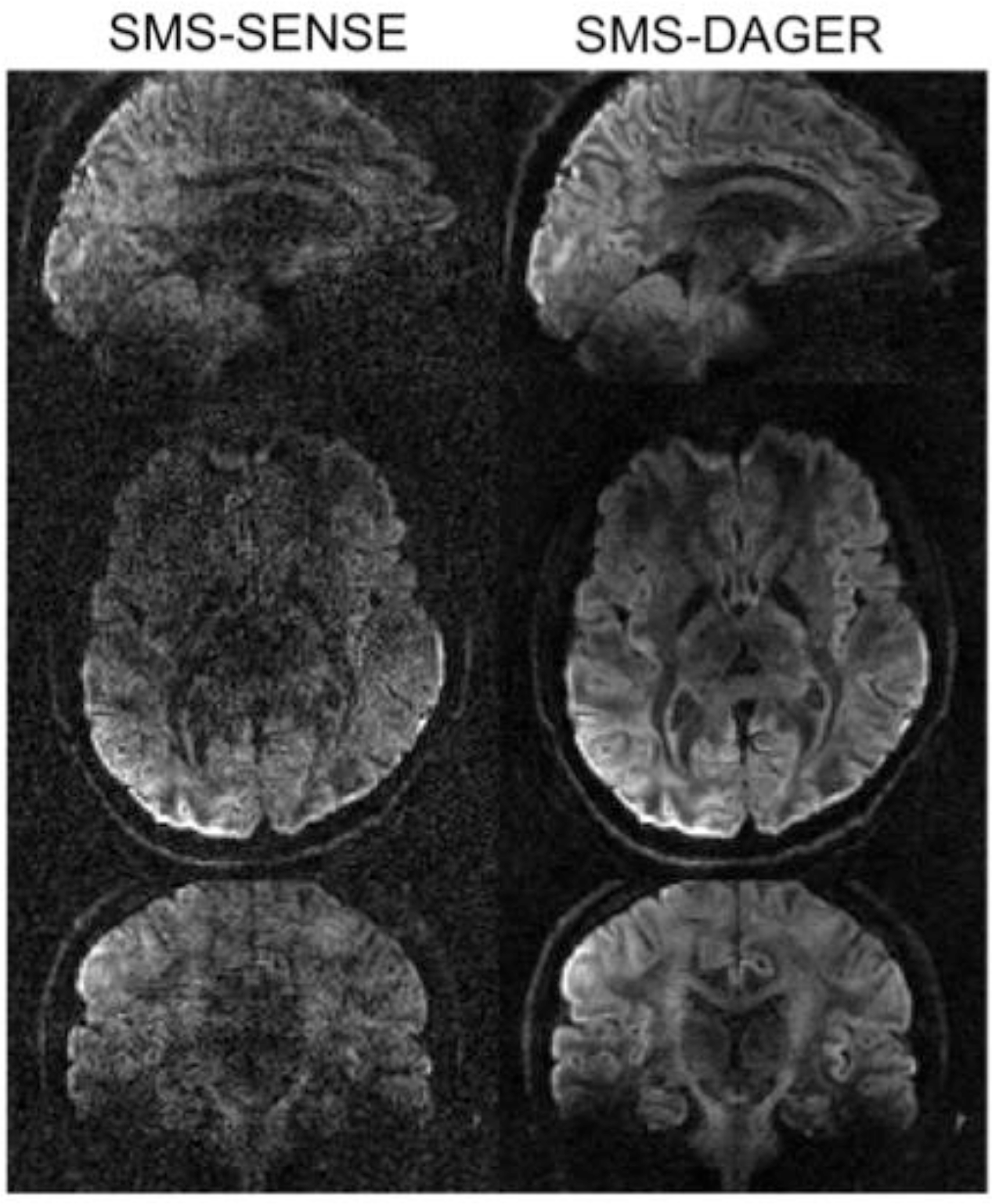
Reconstruction of the same in vivo SMS data using SMS-DAGER and SMS-SENSE, with total acceleration factor 12 (MB=4 and R=3).

Diffusion tensor fits to SMS data in subject 4 are shown in Fig. 6. Tensor metrics derived from SMS-DAGER reconstruction are consistent with the high-SNR SB-3ave reference, while the conventional SENSE reconstruction of the same data results in a noisy estimation of FA and inflated MD values, likely due to the low image SNR and residual aliasing artifacts.

**Figure 6.**
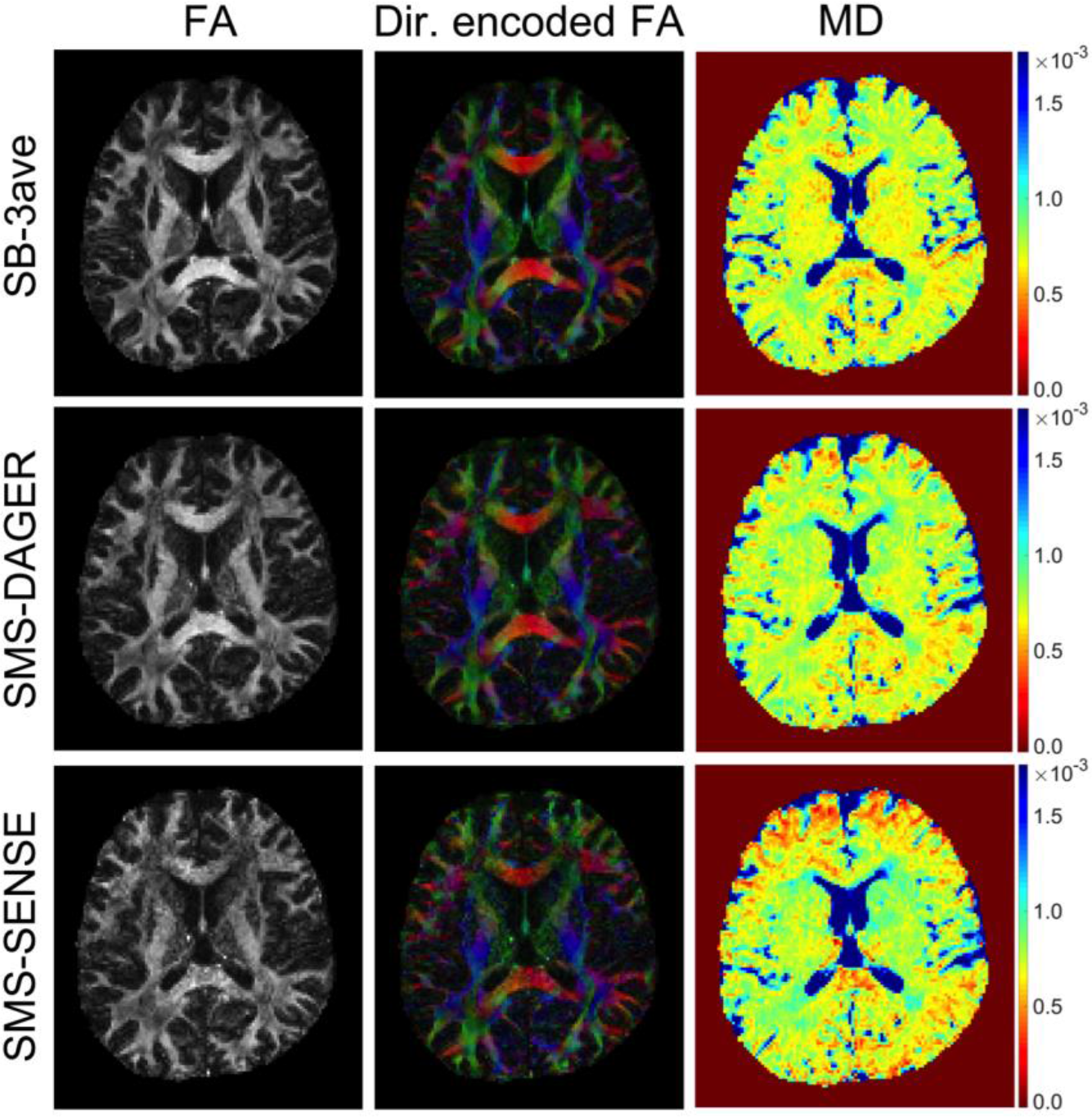
Diffusion tensor analysis of subject 4 data. Comparison of SMS data (MB=4 and R=3) reconstructed using both SENSE and DAGER, and a high-SNR single-band reference (SB-3ave). Three parametric maps are shown here: fractional anisotropy (FA), direction-encoded colormap (red: right-left; green: anterior-posterior; blue: superior-inferior) and mean diffusivity (MD).

### 4.5 Evaluation of angular resolution

If the level of intrinsic signal smoothness is overestimated, DAGER has the potential to produce high-SNR images at the cost of loss of angular information (i.e. smoothing in q-space). In Fig. 7, we investigate the effects of SMS-DAGER reconstruction on angular resolution in comparison to the SB-3ave reference acquisition (non-SMS, with longer scan time). The covariance of the dMRI signal between two images is plotted as a function of the angle between them in q-space (points with angular distance of 0° are the signal variance of a single volume). Angular smoothing would lead to increased covariance at large angular difference. Compared to the high-SNR reference data, SMS images reconstructed with DAGER have only slightly higher covariance, indicating preservation of angular resolution in the DAGER reconstruction. With conventional SENSE reconstruction of the same SMS data, the images are corrupted by noise, resulting in a low covariance between diffusion directions.

**Figure 7.**
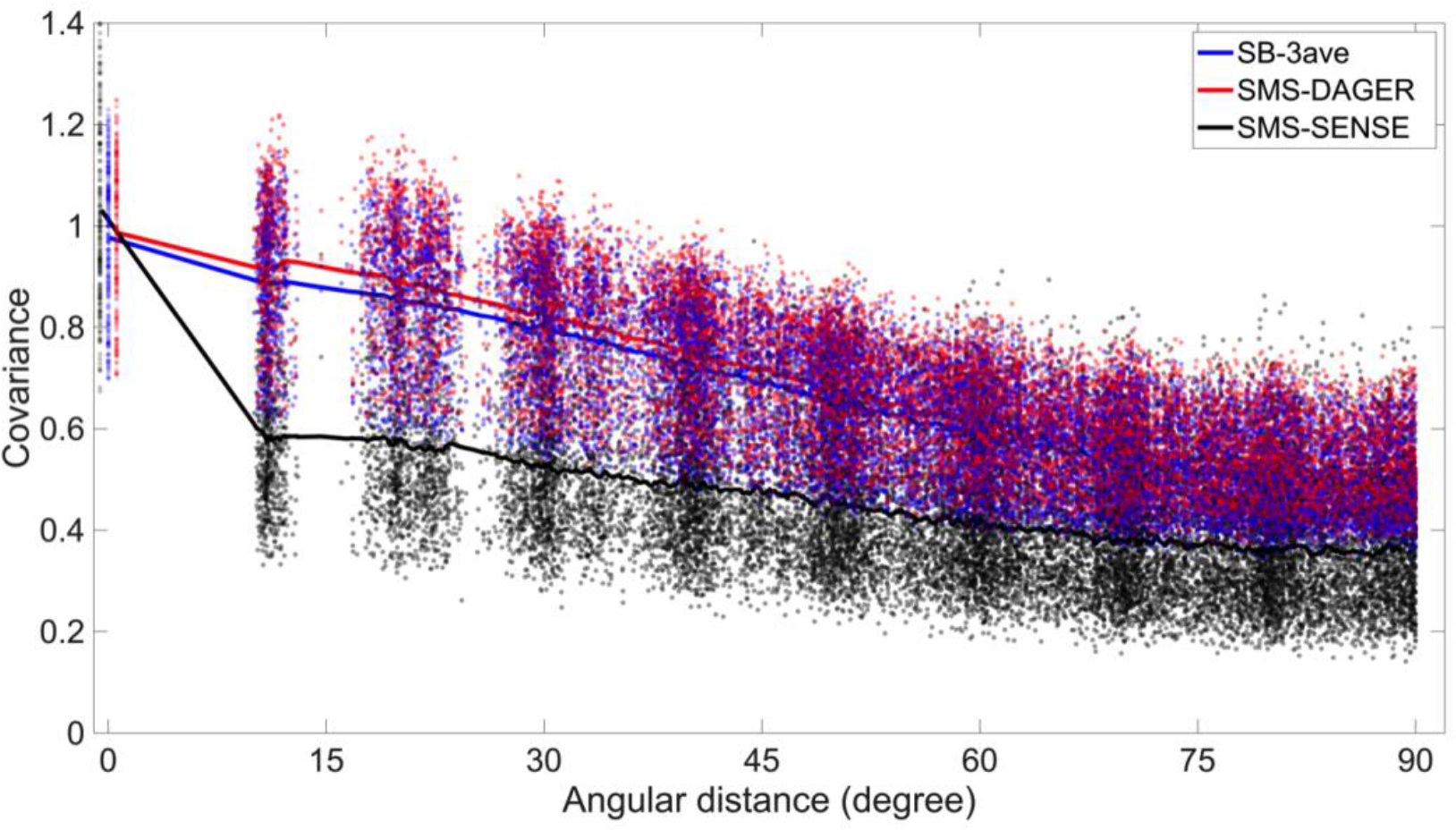
Signal covariance of dMRI images for high-SNR single-band reference (SB-3ave) and SMS data (MB=4, R=3) reconstructed using SMS-DAGER and SMS-SENSE. Each point represents one diffusion direction. The averaged covariances are plotted as solid lines. For each data set, the covariance is normalized by the median of signal variances (angular distance=0°). Note that in the plot we deliberately shift signal-variance points by a small amount along the angular-distance axis for better visualization of the differences.

Comparison of fiber orientations, i.e. the deviation from that estimated from the SB-2ave data, is shown in Fig. 8. For a large number of diffusion directions (128+), the angular differences derived from the SMS-DAGER images are comparable to those derived from the SB-1ave images, with slightly worse performance for the first fiber and equivalent performance for the second fiber (*p* > 0.01, see Supplementary table 1 for details). This suggests preservation of SNR and angular resolution in DAGER reconstruction. The fiber orientations derived from the SMS-SENSE data deviate significantly from the reference, even with many diffusion directions, which is likely due to the noise amplification and residual artifacts in the images.

**Figure 8.**
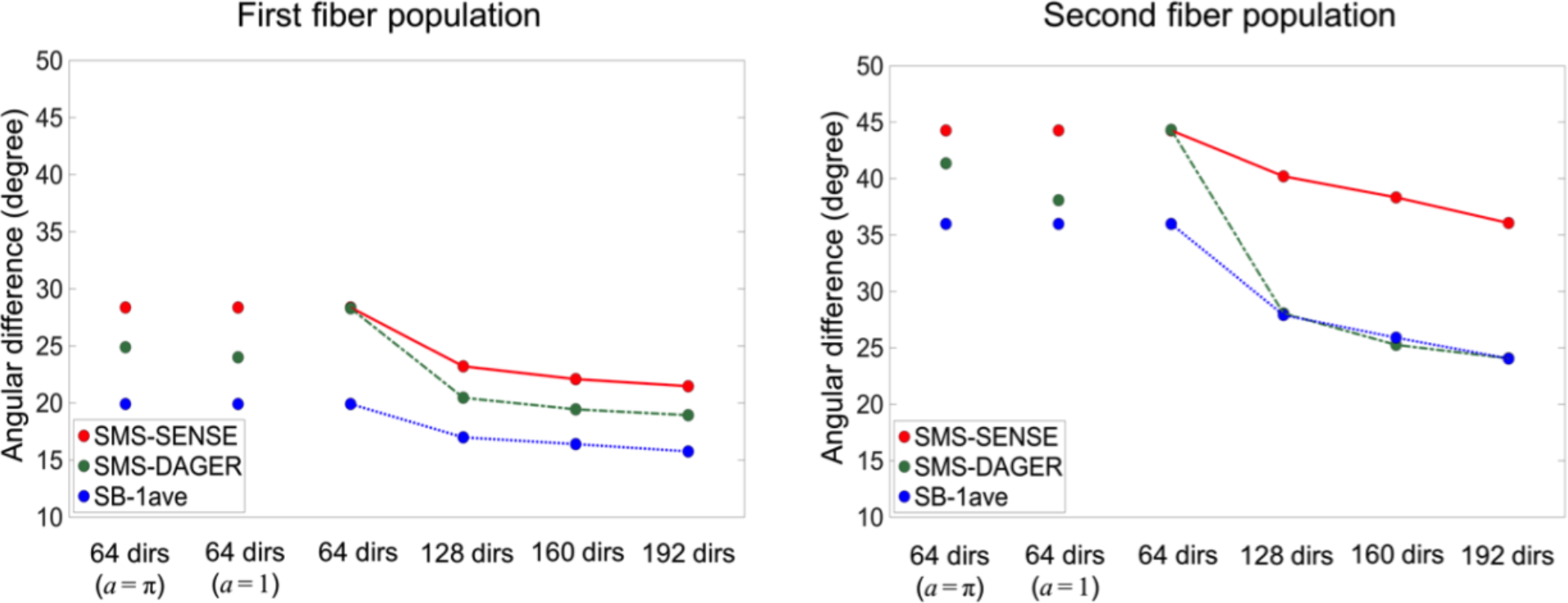
Evaluation of fiber orientations in subject 4, using two averages from the single-band data (SB-2ave) as a reference. The angular difference to SB-2ave is estimated for SB-1ave (the third SB dataset in this subject), SMS-DAGER and SMS-SENSE. Angular estimates are separately considered for the first (left) and second (right) fiber population. Four diffusion direction sets are evaluated, which contain 64, 128, 160 and 192 directions, respectively. For the 64-direction data set, two additional DAGER reconstructions with enforced smoothness (a = 1 and a = π) are also applied (The three 64-direction estimates are identical for SMS-SENSE, and for SB1-ave). For clear visualization of the differences between methods, error bars are not included here.

The number of diffusion directions is expected to be a key determinant of the quality of fiber orientation estimation. On one hand, it limits the number of diffusion volumes that are available to our local GP estimation; on the other hand, it constrains the angular resolution that can be resolved using dMRI. With only 64 directions, SMS-DAGER shows similar performance as SMS-SENSE, presumably reflecting that q-space samples are too far apart to provide useful information for local GP estimation. We can explore the effect of the smoothness hyper-parameter a for this 64-direction dataset, where the DAGER reconstruction does not provide much improvement over conventional SMS-SENSE. If we fix a to an inflated value, the images have higher SNR, suggesting improved conditioning of the reconstruction (see Supplementary Fig. S6). However, this enforced-smoothness constraint may be reasonably hypothesized to introduce angular blurring beyond the intrinsic smoothness of the signal. As shown in Fig. 8, with a slight smoothness constraint (*a* = 1), the estimation of angular resolution is improved. However, when a strong smoothness constraint (*a* = π) is applied, the angular difference is inflated, suggesting a loss of angular resolution in the reconstruction. Indeed, it is noticeable that the image contrast for a single image with *a* = π (indicative of shared information over the entire q sphere) is very similar to a mean diffusivity image (Supplementary Fig. S6).

### 4.6 Diffusion tractography

We conducted automatic mapping of 14 major white matter tracts using AutoPtx (48) on the full-FOV data sets in subjects 1-3. To match the scan time (~11min), SMS-data were acquired with 192 directions and SB data were acquired with 50 directions. The SB data have high SNR due to the long TR required to match the number of slices (12s). As shown in Fig. 9, all of the fiber bundles are robustly identified from the SMS-DAGER data. The single-band results capture many fiber bundles, but its performance is compromised by the limited number of diffusion directions achievable in this scan time. Tracts are in general less abundant in the single-band data and the SMS-SENSE data than the SMS-DAGER data. In some cases, tracts are totally missing at the threshold used in rendering. For example, the acoustic radiation tracts (Fig. 9, middle column, in purple) are clearly captured by the SMS-DAGER data, but essentially absent in both the single-band results and the SMS-SENSE results. Note that these renderings used the same threshold across the different datasets but required these to be determined manually. However, where tracts are largely absent, there was no threshold at which decent tract reconstructions could be produced, and the overall conclusions about the relative performance of the datasets does not change with threshold level.

**Figure 9.**
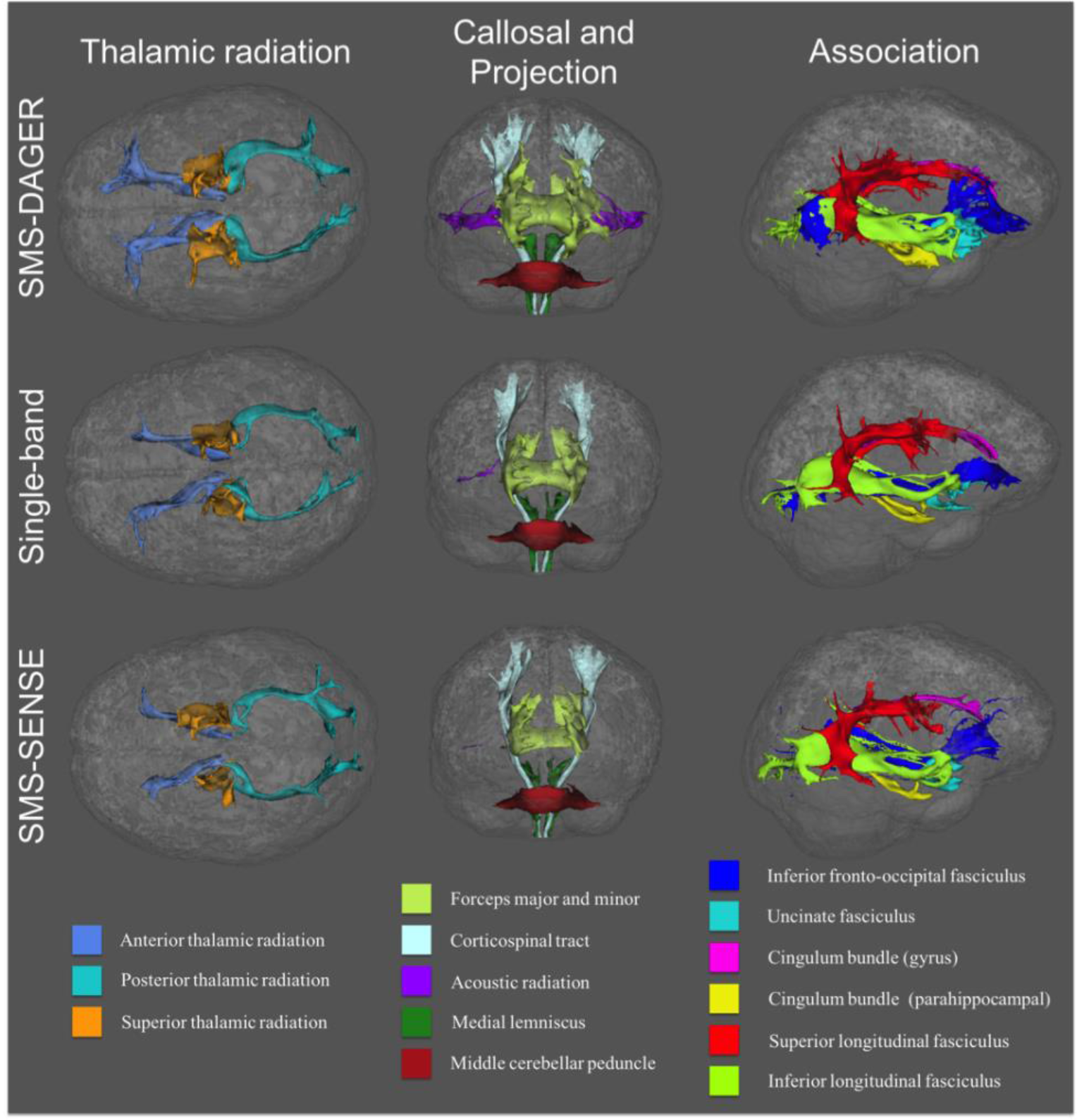
White matter fiber tracts generated using AutoPtx. 14 major white matter pathways (constituting 27 separate tracts in right and left hemispheres) are rendered in superior (left, thalamic radiation), anterior (middle, projection and callosal fibers) and lateral (right, association fibers) views. To match the scan time, SMS data and single-band data contain 192 and 50 directions, respectively. SMS data reconstructed with DAGER support the delineation of all tracts, whereas the single-band data and the SMS data reconstructed with SENSE fail to capture some tracts. Most notable are the acoustic radiations (missing from both SB and SMS-SENSE reconstructions).

## Discussion

This work aims to address one of the major limitations for SMS acceleration: the ill-conditioning of reconstruction at high acceleration factor, which usually results in a strong noise penalty. Improved sampling schemes such as CAIPIRINHA (49) can mitigate this issue, but it remains challenging for diffusion MRI due to its intrinsically low SNR, particularly in combination with in-plane acceleration to reduce distortion and blurring. For example, this limitation was specifically noted in the design of the Human Connectome Project acquisition at 3T, where in-plane acceleration was abandoned due to significant SMS image degradation (33). DAGER is a new approach to improving SMS through a k-q reconstruction that leverages the redundancy of spatial information in q-space. With a maximum scan time, DAGER will allow one to achieve higher acceleration (including higher SMS factor) and thus get more q-space samples (which in turn increase the information available to GP). In comparison to previous work that accelerates purely in q-space (e.g. compressed sensing), k-q acceleration enables improved data fidelity (reduced distortion/blurring with in-plane acceleration) and higher SNR-efficiency (where optimal TR is 1-3 s, depending on field strength (50). Using the DAGER approach, we have demonstrated highly accelerated imaging with superior performance compared with conventional parallel imaging, achieving high SMS acceleration (MB=4) with in-plane undersampling (R=3). Alternatively, DAGER can be used to increase acceleration in one dimension: this was demonstrated here with high in-plane accelerations, but may also enable more ambitious SMS accelerations (higher MB factors).

A unique feature of DAGER is the embedding of Gaussian processes into the k-q reconstruction. Diffusion in biological tissue ranges from free to restricted diffusion, causing the observed diffusion signal in q-space to be relatively smooth. This property has been exploited in Gaussian processes for data analysis; for example, to replace corrupted outlier images with a prediction from adjacent q-space samples (23,25). Unlike methods used in other k-q reconstructions (19,21) that aim to find a parametric model for the diffusion function, Gaussian process method aim to find a parametric model for the covariance function, with the hyper-parameters related to the signal smoothness in q-space learned from the data itself (indeed, GPs can be viewed as a type of machine learning (22)). The optimal hyper-parameters could depend on tissue type, suggesting that it might be preferable to estimate them separately for e.g. gray and white matter. Such a situation would add practical complication to DAGER due to the need for tissue-type segmentation. Fortunately, previous investigations of this found no evidence for overfitting of this kind (24). In addition, the GP approach provides an elegant way to model multi-shell dMRI data, allowing the estimation of data across shells as well as directions (24). Future work will expand the DAGER approach to multiple q-space shells, enabling more sophisticated analyses for microstructural information, such as NODDI (51). Finally, the consistent use of GPs in both data reconstruction and analysis raises the possibility of integrating image reconstruction and data pre-processing, a synergy which could further improve dMRI results.

As DAGER pulls information from adjacent q-space locations, it could introduce angular smoothing if the hyper-parameters overestimate the actual signal smoothness. A direct demonstration of this is given in Fig. 8, where a strong, fixed smoothness constraint led to the loss of angular resolution. We conducted investigations of the effects of DAGER reconstruction on angular resolution. First, we compared the covariance of reconstructed images with respect to angular distance in q-space, finding that the signal covariance derived from the DAGER result is consistent with that from high-SNR reference data (Fig. 7), suggesting that the estimated hyper-parameters are able to capture the underlying covariance of in vivo dMRI signal. Second, we investigated the effects of DAGER reconstruction on voxel-wise fiber orientations in comparison to a reference. For more than 128 directions, angular differences between DAGER and the high-SNR reference are not considerably larger than those derived from the conventional single-band data (Fig. 8). However, with a smaller number of directions, DAGER provided considerably less benefit. These evaluations and investigations suggest DAGER reconstruction is able to provide a good preservation of angular resolution.

The effects of DAGER on automated tractography are studied in data sets with matched scan time. Tractography using the DAGER reconstructions of SMS data out-performed SENSE reconstruction of the same data, as well as single-band data with matched scan time. DAGER produced a more extensive and comprehensive representation of all tracts considered, particularly those involving crossing fiber regions. These results demonstrate improvements of DAGER in a domain that is important in practice (beyond producing high-quality images).

As DAGER relies on the joint information between q-space neighbours, the number of diffusion directions is expected to affect the reconstruction performance. In this work, we found that DAGER reconstruction with 64 diffusion directions is very close to SENSE reconstruction, indicating low covariance between the available q-space samples (the average angular difference between each point and its nearest neighbour is about 20 degree). This result suggests that a relatively large number of directions are needed for DAGER reconstruction. However, given that DAGER is intended to support higher SMS acceleration factors, the burden of time required to reach a target number of directions goes down, leading to a synergy between the needs of the GP and the benefits it delivers. Here, we were able to achieve high quality reconstructions by acquiring 128 directions in ~7.5min. Additionally, once distortion correction is incorporated in DAGER, we may be able to exploit q-space symmetry (the equivalence of q-space samples on opposite sides of the sphere) to increase the number of ‘effective’ q-space neighbours. This will be investigated in future work.

The hyper-parameters in the GP model are estimated from the data based on a Bayesian formulation. One subtle but important detail is that image aliasing, which the reconstruction aims to remove based on smoothness, has a detrimental effect on hyper-parameter estimation. An iterative updating scheme is adopted in DAGER, where the hyper-parameter estimates and the image reconstruction are consecutively updated such that errors in both procedures are gradually suppressed over multiple iterations.

It is worth noting that this reconstruction scheme differs from classical model fitting procedure where the model is identical in each iteration. Instead, in DAGER reconstruction the model is refined in each iteration, which might lead to a non-monotonically decreasing cost-function. In our experiments, we find the reconstruction typically converges after 10-15 iterations and the model becomes relatively stable afterwards. This is demonstrated in simulation (Fig. 2), where the iteration converges after 13 iterations. However, under certain circumstances, the proposed algorithm might fail to converge, such as strong subject motion, severe eddy current distortion and a low number of diffusion directions. In future work, we aim to integrate the motion and distortion correction, which could aid in convergence under these conditions.

The calibration of noise variance can affect the reconstruction performance, especially for low SNR data (see Supplementary Fig. S3), which are common in diffusion MRI. In simulation, DAGER’s noise variance estimates were demonstrated to be close to the ground truth, but even more importantly the image errors (NRMSE) using the estimated noise variance were very similar to that of a reconstruction with the correct noise variance. In addition to providing high quality reconstructions, the automatic determination of reconstruction parameters is pragmatic, as no manual adjustment is needed, unlike many previous k-q methods.

For this proof-of-principle study, we focus on the reconstruction of dMRI data and acquire b=0 data using a multi-shot EPI protocol that samples full k-space to avoid strong reconstruction artifacts at high acceleration. However, the multi-shot sequence will increase the sensitivity to subject motion if the acquisition time is too long, even for b=0 data without significant shot-to-shot phase errors. This issue could be mitigated by sampling partial k space (reducing the number of segments), using lower slice acceleration (e.g. MB=2) and/or acquiring single-band b=0 data.

Similar to other parallel imaging reconstruction techniques, performance of DAGER is affected by multiple factors. We investigated the effect of the number of diffusion directions at a given acceleration, which suggests a relatively large number of directions are required for a 12-fold acceleration (i.e., MB=4, R=3) using DAGER. Similarly, for a fixed number of diffusion directions we expect a limitation on the acceleration factor that can be achieved. However, the achievable acceleration factor will depend on a number of additional factors, including receive coil, field strength, b value, RF pulses, k-q sampling and spatial resolution. It is also expected that different analysis methods and models will vary in their sensitivity to noise amplification and angular smoothness. As such, it is not straightforward to determine an appropriate acceleration factor without resorting to some empirically determined heuristics that likely need to be based on the final outputs of the target data analysis.

Another limit to acceleration with DAGER relates to the need to estimate and account for motion-induced phase errors. Our navigator-free approach uses conventional parallel imaging methods to estimate phase errors (31). This approach would fail at acceleration levels where parallel imaging is not able to provide an accurate image reconstruction. Advanced reconstruction methods, such as low-rank modelling and compressed sensing, may be able to support higher acceleration factors. Alternatively, DAGER can use explicitly-acquired navigators. At ultra-high field, where SMS acceleration is generally limited by SAR, the need for an additional refocusing pulse for a navigator is problematic. At lower field, acquisition of navigators is more feasible, incurring a slight loss of efficiency due to longer TR.

Although the current work only demonstrates accelerated 2D and 2D-SMS dMRI, other acquisition methods like 3D multi-slab should also be compatible with DAGER provided an incoherent k-q sampling scheme is incorporated. Recently, several studies have demonstrated the use of simultaneous multi-slab acquisition for dMRI, which enables optimal SNR efficiency with shorter scan time (52,53). It is expected that the combination of DAGER and simultaneous multi-slab acquisition can achieve even higher scan efficiency with improved image quality.

Finally, parallel imaging reconstruction is a mature field, with a broad range of techniques from SENSE to GRAPPA (4,5,41,42,54) and many approaches to regularization (6–10). All of these techniques can potentially be embedded within the DAGER framework, and thus have potential to improve DAGER, particularly at the limits of the current implementation of DAGER (e.g., at high acceleration or with a reduced number of diffusion directions). However, the primary goal of this paper is to demonstrate that sharable information in q-space can be leveraged to improve upon existing parallel imaging reconstructions. Exploration of how well different regularization techniques are able to take advantage of this shared information will be the topic of future work.

## 5. Conclusion

In this work, we developed a method to accelerate dMRI acquisition, which incorporates sharable information between q-space samples to improve the reconstruction of k-space data. This k-q reconstruction approach uses Gaussian processes to estimate and exploit local smoothness without introducing undesirable angular blurring. Results from simulation and in vivo data demonstrate the efficacy of the proposed method, particularly for high SMS acceleration with in-plane undersampling.

## 6. Acknowledgement

We wish to thank Cornelius Eichner and Kawin Setsompop for sharing their MATLAB-based MultiPINS RF pulse design tool. This work is supported by the Wellcome Trust (202788/Z/16/Z, KM and 098369/Z/12/Z, JA), Marie Curie Initial Training Network Program (FP7-PEOPLE-2012-ITN-316716, WW). The Wellcome Centre for Integrative Neuroimaging is supported by core funding from the Wellcome Trust (203139/Z/16/Z).

